# Directed cell-type recruitment during consolidation of a remote memory

**DOI:** 10.1101/2025.06.13.659562

**Authors:** Zachary Zeidler, Abigail L. Yu, Moonkyu Patrick Seong, Rio Hundley, Laura A. DeNardo

## Abstract

Memories are consolidated into a distributed neocortical network for long-term storage. Long-term memory retrieval relies on cells that are active during learning and undergo necessary plasticity. However, remote memory retrieval activates a broader circuit, with learning-activated cells representing only a small subset. What are the rules and cell-types governing memory trace reorganization? We identified a class of prefrontal projection neurons that are gradually recruited to a memory trace through synaptic activity of learning-activated cells. This population, which projects to the temporal association area (TEa), progressively strengthens its encoding of memory-induced behaviors, mirroring increases in TEa activity. Notably, the prefrontal-TEa pathway is required for remote but not recent memory retrieval. Our findings reveal a cell type-specific mechanism underlying memory trace reorganization during consolidation.

## Main

While the processes underlying memory encoding and retrieval of recent memories have been extensively studied^1–7^, the cellular and circuit mechanisms necessary to establish a remote memory remain elusive^8^. Current theories of memory consolidation suggest that enduring memory traces are gradually encoded in distributed neocortical networks^8–11^. Yet the rules of how these neocortical memory networks are built remain unclear. Understanding the principles governing the construction of these networks, as well as the synaptic architecture of neocortical memory networks, will provide key insights into mechanisms of memory consolidation.

The medial prefrontal cortex (mPFC) is critical for consolidating long-term memories^12^ but the specific cell types and circuits involved remain poorly understood. Neurons in the mPFC that are active during memory encoding undergo lasting synaptic changes essential for remote memory retrieval^13^. These changes, including spine growth and strengthened excitatory synaptic connectivity, make those neurons more likely to be reactivated by memory cues, facilitating remote memory recall^13,14^. However, the neurons activated during learning represent only a subset of those engaged during remote memory recall^13–15^, and distinct mPFC neurons^15^ and efferent pathways^16^ contribute to memory retrieval across time. Yet the identity, connectivity, and behavioral functions of the mPFC neurons that are added to the remote memory trace remain poorly understood.

mPFC contains many classes of projection neurons that have distinct functions^15,17,18^. Of these, intratelencephalic (IT) neurons, which project to other cortical areas*(17)*, are well positioned to coordinate the formation of distributed neocortical memory networks. One major target of mPFC IT neurons is the temporal association cortex (TEa)^19,20^, which encodes conditioned stimuli and relays this information to the amygdala^21^. While TEa itself plays a specialized role in remote memory^22,23^, the role of the mPFC-TEa pathway is unknown. We hypothesized that mPFC-TEa neurons may be recruited to a memory trace across time, forming a neocortical network that mediates long-term memory.

## mPFC-TEa neurons gain encoding of conditioned cues and behavior across time

To investigate how mPFC-TEa projection neurons encode memory-relevant features across time, we recorded Ca^2+^ activity in these cells *in vivo* during auditory fear conditioning (FC) and during recent (1 day; D1) and remote (28 day; D28) memory retrieval (Fig. 1A). We compared mPFC-TEa neurons to the general population of mPFC neurons (Fig. 1B). There was no difference in freezing, a proxy of memory strength, between these groups during memory testing (Fig. 1C). To determine whether single neurons selectively encode memory information, we computed a receiver operating characteristic (ROC) curve—which quantifies stimulus detection strength over a range of binary thresholds—for each neuron’s response to the conditioned tone and freezing behavior (Extended Data Fig. 1A). From the recent to the remote timepoint, we observed a significant increase in the number of mPFC-TEa neurons that encoded the conditioned tone (Fig. 1D), whereas the proportion of mPFC-TEa neurons that were modulated by freezing behavior was stable across time (Fig. 1E). This effect was driven by an increase in tone-excited rather than tone-suppressed cells (Extended Data Fig. 1B-C). The targeted increase in tone-modulated neurons suggests enhanced representation of memory cues and behavior linked to those cues, rather than a generalized increase in activity or enhanced representations of fearful behaviors. In contrast, the general mPFC population exhibited similar proportions of neurons responding to the conditioned tone and/or freezing behavior across time (Fig. 1D-F; Extended Data Fig. 1D,E). Together, these data suggest that while a stable number of mPFC neurons encode the conditioned tone, mPFC-TEa neurons make up a larger proportion of this population across time.

**Figure 1.**
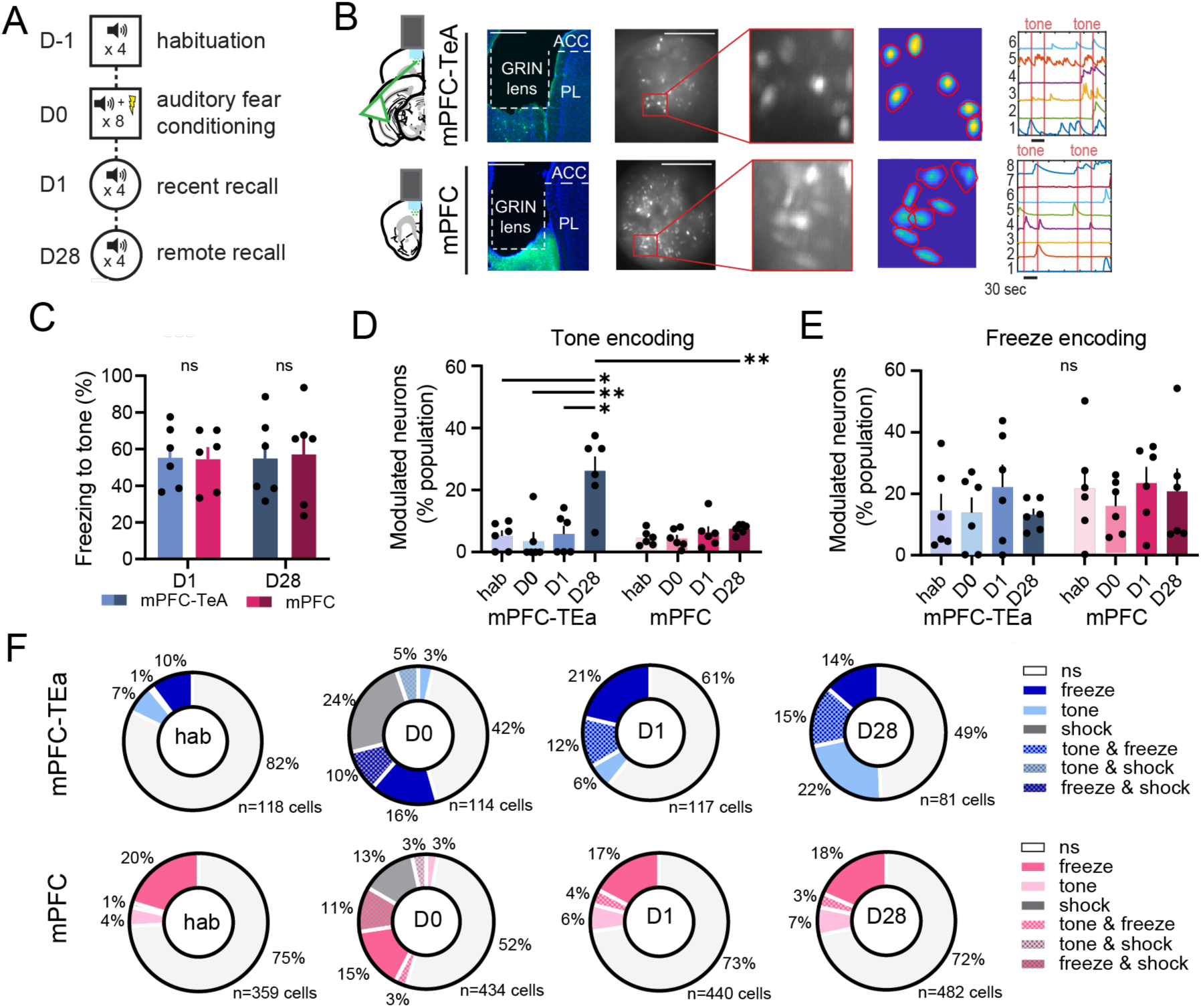
– Representations of memory features gradually accumulate in mPFC-TEa neurons. **A.** Cartoon of experimental timeline while recording neuronal activity. **B**. Example miniscope recordings showing lens acement, total field of view, identified ROIs, and extracted calcium traces in mPFC-TEa neurons (top) and general PFC neurons (bottom). **C**. Tone freezing for all mPFC and mPFC-TEa groups across time (Group F(1,10)=0.7, P=0.4; me F(1,10)=0.3, P=0.5, Group x Time F(1,10)=0.01, p = 0.9; Two-way Repeated Measures ANOVA, n=6 mice per oup). **D**. Proportion of tone encoding neurons in mPFC-TEa and mPFC recordings across time (Group F(1, 10)=9.6, =0.01, Time F(2.4, 24.7)=11.3, P=0.002, Group x Time F(3, 30)=6, P=0.001; Two-way Repeated Measures ANOVA, ukey’s multiple comparisons tests, n=6 mice per group). **E**. Proportion of freeze encoding neurons in mPFC-TEa and PFC recordings across time (Group F(1, 10)=0.7, P=0.4, Time F(2.2, 22)=0.9, P=0.4, Group x Time F(3, 30)=0.2, =0.8, Two-way Repeated Measures ANOVA, n=6 mice per group). **F**. Pie charts showing functional breakdown of PFC-TEa (top) and mPFC (bottom) recordings across time. <0.05, **P<0.01. Error bars represent mean ± s.e.m.

To begin to understand how mPFC-TEa neurons may be recruited into the remote memory trace, we tracked individual mPFC-TEa neurons across imaging sessions and examined their activity during memory encoding and recent and remote memory retrieval (Extended Data Fig. 1F). We focused on the substantial proportion of mPFC-TEa neurons that encoded the aversive foot shock during FC (Fig. 1G). While some reliably increased their activity during foot shocks, others reliably decreased their activity (Extended Data Fig. 1G). Shock-sensitive mPFC-TEa neurons did not preferentially encode the conditioned tone during recent memory retrieval (Extended Data Fig. 1H). However, during remote memory retrieval, the same neurons encoded the tone at rates significantly greater than would be predicted by chance (Extended Data Fig. 1H). Interestingly, shock-suppressed neurons were more likely to become tone-excited neurons than their shock-excited counterparts, which were predominantly unresponsive or suppressed by the tone during remote recall (Extended Data Fig. 1I). This suggests that a population of shock-suppressed mPFC-TEa neurons remains relatively silent at recent timepoints, and then progressively integrates into the remote memory trace, gaining tone-responsivity across time.

## Functional specialization of mPFC-TEa population activity emerges over time

The mPFC contains functionally heterogenous subclasses of projection neurons, and subsets of mPFC neurons can encode distinct aspects of conditioned stimuli (e.g. onset, offset, or intersections with behavior) ^18,19,24–26^. To better understand the functional profiles of mPFC-TEa neurons that are incorporated into the remote memory trace, we investigated whether mPFC-TEa neurons encode particular aspects of the conditioned tone across time, and whether they represent a functionally-defined subclass of mPFC neurons. To do so, we pooled together all neurons from all mPFC and mPFC-TEa recordings and performed hierarchical clustering based on their activity patterns during the tone (Fig. 2A, Extended Data Fig. 2A,B). This approach groups together neurons with similar tone responses and allows a comparison between mPFC-TEa neurons and the general mPFC population.

**Figure 2.**
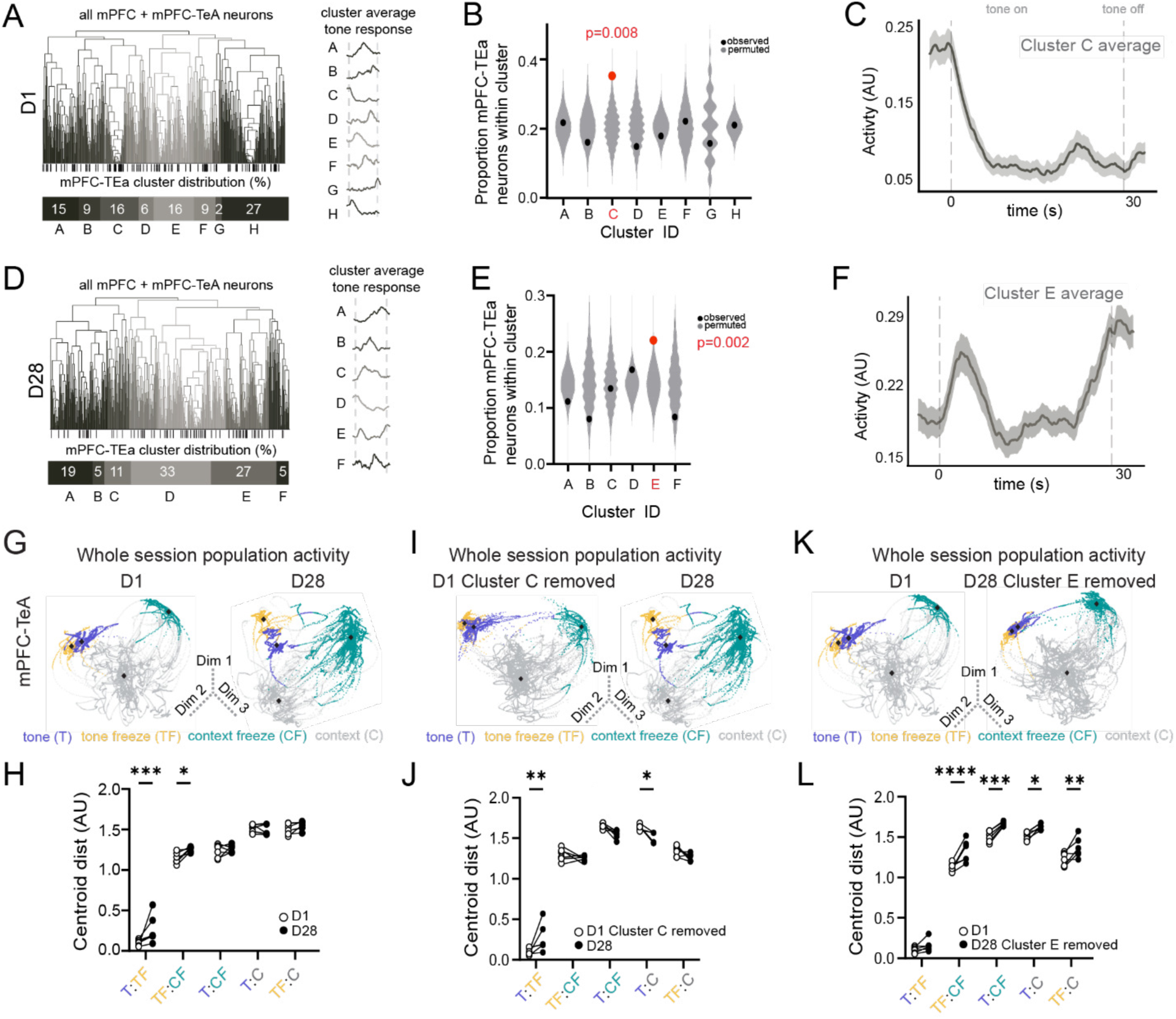
– Functional specialization of mPFC-TEa population activity emerges over time. **A.** Left: clustering of all mPFC and mPFC-TEa tone responses on D1. Shade indicates different clusters, black lines below indicate mPFC-TEa neurons. Bar graph shows percent of total mPFC-TEa neurons assigned to that cluster. Right: cluster average tone response. **B.** Permuted (gray) and observed (black, red) proportion of mPFC-TEa neurons in each cluster luster A: P=0.45; Cluster B: P=0.21; Cluster C: P=0.008; Cluster D: P=0.2; Cluster E P=0.26; Cluster F: P=0.45; Cluster G: P=0.42; Cluster H: P=0.49). **C.** Cluster C averaged tone response. **D.** Same as A for D28. **E.** Same as B for 28 clusters (Cluster A: P=0.13; Cluster B: P=0.12; Cluster C: P=0.49; Cluster D: P=0.18; Cluster E: P=0.002; Cluster F: =0.14). **F.** Cluster E averaged tone response. **G.** Example embeddings of data from an mPFC-TEa recording on Day 1 op) and Day 28 (bottom. Each point corresponds to one frame, color coded by behavior. Black dot indicates centroid of behavior cluster. **H.** Distance between centroids at D1 vs. D28 for mPFC-TEa recordings (Centroids F(4,25)=220.1, <0.0001, Time F(1,25)=8.34, P=0.007, Centroids x Time F(4,25)=3.5, P=0.01, Two-way Repeated Measures ANOVA, dák’s multiple comparisons test, n=6 mice). **I.** Same as G after removing neurons belonging to Cluster C on Day 1.J. me as H after removing Day 1 Cluster C neurons (Centroids F(4,25)=793, P<0.0001; Time F(1,25)=3.1, P=0.08; entroids x Time F(4,25)=7, P=0.0006, Two-way Repeated Measures ANOVA, Šídák’s multiple comparisons tests, n=6 ice). **K.** Same as I after removing neurons belonging to Cluster E on Day 28. **L.** Same as H after removing Day 28 Cluster 5 neurons (Centroids F(4,25)=512, P<0.0001; Time F(1,25)=72, P<0.0001, Centroids x Time F(4,25)=2.2, =0.09, Two-way Repeated Measures ANOVA, n=6 mice). <0.05, **P<0.01, ***P<0.001. Error bars represent mean ± s.e.m.

During recent memory retrieval, we identified eight functional clusters with diverse tone-evoked responses (Fig. 2A). While we observed mPFC-TEa neurons in all clusters, Cluster C, containing 16% of all mPFC-TEa neurons, had more mPFC-TEa neurons than expected by chance (Fig. 2B) and exhibited a marked suppression during the tone (Fig. 2C). During remote memory retrieval, we identified six functional clusters, exhibiting a broad range of tone-evoked response profiles (Fig. 2D). mPFC-TEa neurons were again present in all clusters, but Cluster E, containing 27% of all mPFC-TEa neurons, had more mPFC-TEa neurons than expected by chance (Fig. 2E). This cluster exhibited a transient increase in activity following the onset of the tone, followed by late ramping activity through the end of the tone (Fig. 2F). We then tracked which neurons belonged to which cluster at each timepoint. Most mPFC-TEa neurons changed their dynamics between D1 and D28 (Extended Data Fig. 2C). These results show that mPFC-TEa neurons have diverse cue-induced dynamics during memory retrieval, but, as a class, exhibit biases toward particular activity profiles that evolve between recent and remote timepoints.

To understand the relevance of these functional clusters, we examined how they contribute to mPFC population codes. During presentations of aversive cues, mPFC population codes contain high dimensional representations of threat-conditioned behaviors^27–30^. We therefore examined how mPFC-TEa population activity encodes key behavioral states and then isolated the contributions of the functional subpopulations.

We characterized mPFC-TEa population activity using CEBRA, a self-supervised network that learns a low-dimensional embedding of population activity by incorporating guiding labels^31^. Separate networks were trained with tone, freezing, or both as labels (Extended Data Fig. 3A). The networks trained with both tone and freezing labels captured neural data better than tone or freezing alone (Extended Data Fig. 3B). However, data from remote recall outperformed recent recall in the combined tone-freezing condition, indicating that mPFC-TEa population activity is increasingly tuned to memory-guided behavioral activity across time (Extended Data Fig. 3C,D).

We used these embeddings to examine how mPFC population activity represents the tone, freezing, and their intersections. We plotted low-dimensional embeddings of the entire recording sessions, plotting each frame as a point in activity space. Frames were color-coded according to their corresponding behavioral state: tone-induced freezing, contextual freezing, tone without freezing, or context (i.e. no tone) without freezing. To quantify how distinctly mPFC population activity represents these states, we measured the distances between the centroids of each behavioral state (Fig. 2G). The centroids for tone and tone-induced freezing were further apart on D28 compared to D1, as were the centroids for tone-induced freezing and contextual freezing (Fig. 2H). This increased distance in neural state-space indicates that the population-level representations of cued freezing in mPFC-TEa neurons become increasingly distinct from their representations of the cue across time. Notably, no such temporal changes were observed in the overall mPFC population activity (Extended Data Fig. 4A,B).

We next investigated whether neurons in the specialized clusters drove these changes in ensemble activity. To test this, we performed *in silico* lesions by removing neurons from mPFC-TEa enriched clusters and regenerating the embeddings. On Day 1, removing Cluster C neurons did not alter the separation of tone and tone-freezing clusters (Fig. 2I,J). However, on Day 28, removing Cluster E eliminated the separation between tone and tone-freezing in population state space (Fig. 2K,L). This effect suggests that Cluster E contributes to population coding by distinguishing between memory cues and memory-guided behaviors. The specificity of these effects was confirmed by a control analysis in which equivalent numbers of neurons were removed from random clusters. In this case, the tone and tone-freezing centroids remained separated (Extended Data Fig. 4C,D). Together, these findings indicate that functional subclasses of mPFC-TEa neurons drive time-dependent changes in population activity, ultimately forming a distinct representation of tone-induced freezing during remote memory retrieval.

## The mPFC–TEa pathway supports remote but not recent memory recall

Based on the enhanced representations of the tone and tone-induced freezing in mPFC-TEa neurons during remote recall, we hypothesized that the mPFC-TEa circuit would be important for remote but not recent memory. To investigate this, we virally-delivered an inhibitory opsin^32^ in mPFC neurons and implanted an optic fiber bilaterally over TEa (Fig. 3A). After fear conditioning, we optogenetically inhibited mPFC-TEa axons during either recent or remote memory retrieval in separate groups of mice. Inhibiting mPFC-TEa projections during recent memory retrieval had no effect on freezing behavior, yet it significantly reduced freezing during remote memory retrieval (Fig. 3B). These effects were specific to tone-conditioned freezing, as we observed no effects on contextual freezing (Fig. 3C) or on tone freezing in fluorophore-only control animals (Extended Data Fig. 5A,B). The specificity of this circuit was confirmed by performing a similar experiment on mPFC projections to the basolateral amygdala, which, consistent with previous findings^16^, reduced freezing during recent but not remote memory retrieval (Extended Data Fig. 5C,D). These results show that the mPFC-TEa circuit in has a preferential role in remote cued memory recall.

**Figure 3.**
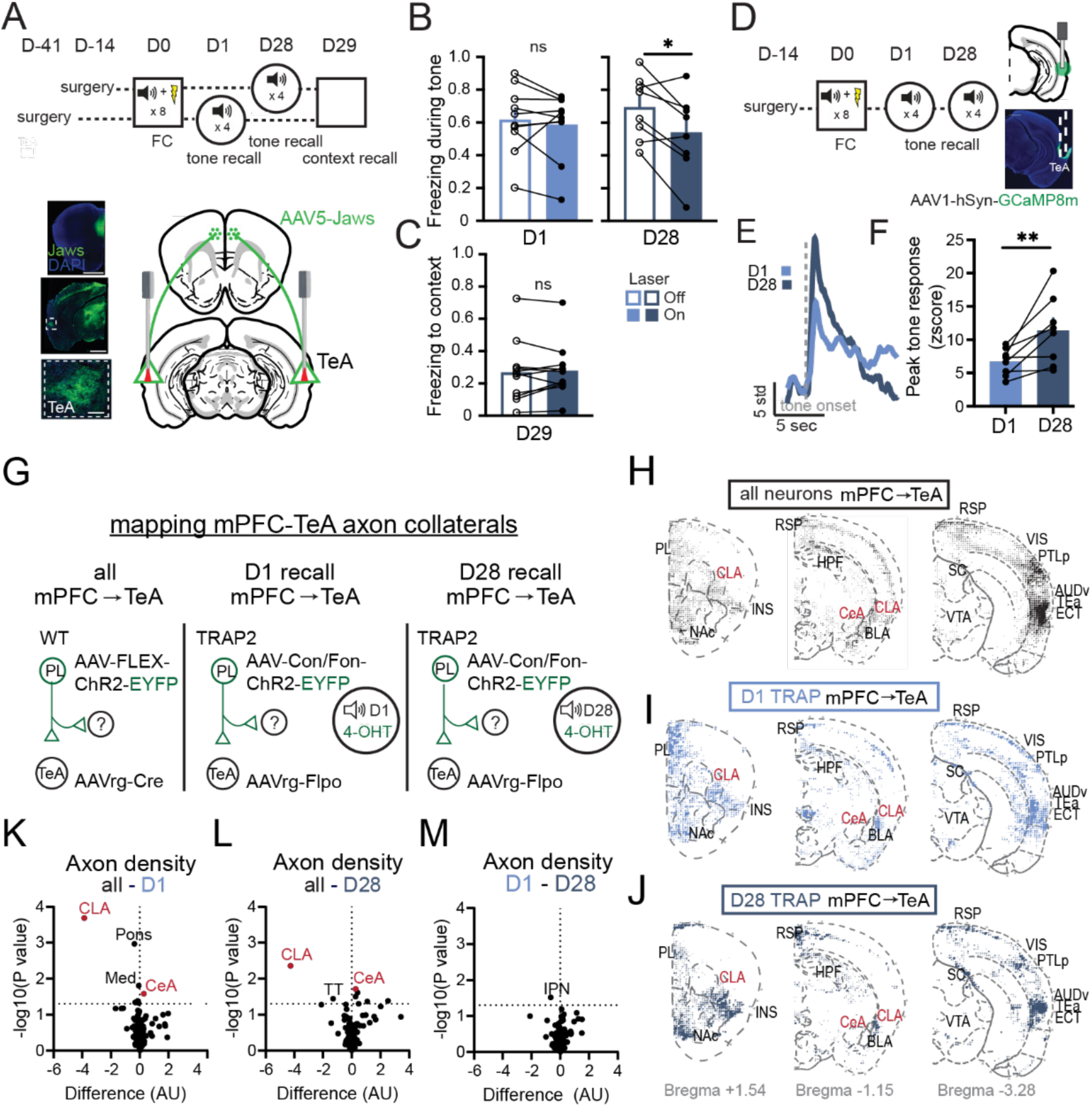
– Time-dependent changes in the behavioral role—but not axon collaterals—of the mPFC–TEa pathway. **A**. Schematic of experimental paradigm for auditory fear conditioning and recall testing (top), and surgery scheme for circuit manipulation (bottom). B. Tone-induced freezing at recent (D1) and remote (D28) recall with and without laser 1 paired t-test=0.4, n=10 mice; D28 paired t-test P=0.02, n=8 mice). **C.** Contextual freezing on D29 in Jaws and GFP mice (paired t-test, P=0.5, n=11 mice). **D.** Schematic for recording TEa bulk activity using fiber photometry during recent 1) and remote (D28) memory recall. **E.** Example traces of TEa activity during tone onset. **F.** TEa tone responses during cent and remote memory retrieval (paired t-test, P=0.01, n=8 mice).**G.** Schematic of surgical scheme to label the axonal ojections of all mPFC-TEa neurons (left), mPFC-TEa neurons active on D1 (middle), or mPFC-TEa neurons active on 28 (right). **H.** Dotogram showing density of axonal innervation across brain segments in general mPFC-TEa neurons. Dot size represents normalized pixel counts summed across subjects (n=5 all mPFC-TEa mice; n=4 D1 mPFC-TEa mice; 5 D28 mPFC-TEa mice). **I.** Same as I for mPFC-TEa neurons active during recent memory retrieval. **J.** Same as I for PFC-TEa neurons active during remote memory retrieval. **K.** Volcano plot comparing regional axonal innervation density between the all mPFC-TEa cohort and recent retrieval (D1) activated mPFC-TEa cohort (CLA difference=-3.861 arbitrary units (AU), –log10(P)=3.688, CeA difference=0.288 AU, –log10(P)=1.584). **L.** Same as L comparing all mPFC-Ea and mPFC-TEa neurons active during remote retrieval (CLA difference=-4.2 AU, –log10(P)=2.359, CeA fference=0.276 AU, –log10(P)=1.713). **M.** Same as L comparing mPFC-TEa neurons active during recent and remote emory retrieval. CLA = claustrum, CeA = central amygdala, IPN = interpeduncular nucleus. <0.05, **P<0.01. Error bars represent mean ± s.e.m.

To better understand the role of the target region, TEa, across time, we used fiber photometry to record bulk Ca^2+^ activity from TEa neurons during recent and remote recall (Fig. 3D). We observed a time-dependent increase in tone-induced fluorescent activity from recent to remote memory recall (Fig. 3E,F). Non-shocked control mice did not exhibit a temporal change in TEa activity levels (Extended Data Fig. 5E,F), indicating that TEa’s pronounced response during remote recall is both time– and learning-dependent.

## Mapping the brain-wide organization of recall-activated mPFC-TEa axonal projections

To clarify how mPFC-TEa neurons support memory retrieval across time, and to learn more about the synaptic organization of memory networks, we examined the brain-wide axonal projections of activity-defined mPFC-TEa neurons. Even among mPFC-TEa neurons, distinct subclasses send axon collaterals to distinct sets of target regions^20^. We therefore mapped the brain-wide projections of mPFC-TEa neurons and investigated whether mPFC-TEa neurons specifically active during recent versus remote memory recall exhibit distinct axonal collateralization patterns, potentially reflecting anatomical subclasses. To test this, we performed viral tracing in Fos^2A-iCreERT2^ (TRAP2) mice^15^, which allowed permanent expression of fluorophores in neurons that were transiently active during FC. We then used tissue clearing and light sheet microscopy to compare the brain-wide axon collateral projections of mPFC-TEa neurons that were active during recent or remote memory retrieval, as well as the general population of mPFC-TEa neurons (i.e., not defined by activity; Fig. 3G-J; Extended Data Fig. 6).

Compared to the general mPFC-TEa population, mPFC-TEa neurons that were active during both recent and remote memory retrieval had more axon collaterals in the claustrum and fewer in the central amygdala (after normalizing to total axon density; Fig. 3K,L). In contrast, we did not observe major differences in the overall projection patterns of mPFC-TEa neurons active during recent versus remote memory (Fig. 3M). These findings indicate that memory-activated mPFC-TEa neurons represent a distinct subclass of the broader mPFC-TEa population that preferentially targets the claustrum. However, mPFC-TEa neurons involved in recent and remote memory project to similar sets of downstream regions. This suggests that changes in the neuronal dynamics—not changes in the anatomical identity—of active mPFC-TEa neurons drive the pronounced shift toward tone-encoding during remote recall.

These data also reveal that beyond TEa, the mPFC-TEa neurons active during memory retrieval send axon collaterals to several additional cortical areas. These include the insular cortex (INS), the retrosplenial cortex (RSP), the ventral auditory area (AUDv), the ectorhinal area (ECT), the posterior parietal cortex (PTLp), and subcortical areas including the claustrum, basolateral amygdala, and nucleus accumbens (NAc) (Fig. 3I,J). These findings provide key insight into the synaptic organization of a distributed cortical network that supports remote memory.

## Synaptic inhibition of mPFC learning-activated cells impairs the formation of a remote memory trace

The formation of a remote memory trace in the mPFC depends upon both local mPFC activity and synaptic input from upstream regions^13–15^. Given the dense recurrent connectivity within mPFC^33–35^—which strengthens during memory consolidation^13^—we hypothesized that synaptic activity from learning-activated mPFC neurons could gradually recruit mPFC-TEa neurons to the remote memory trace. To test this, we blocked synaptic output from learning-activated mPFC neurons beginning in the days following learning and assessed the impact on remote memory. Using TRAP2 mice, we permanently expressed tetanus toxin light chain (TeNT), which blocks synaptic vesicle fusion and therefore synaptic transmission^36^, in mPFC neurons that were active during FC. In the same mice, we fluorescently labeled mPFC-TEa neurons using an axon-transducing AAV (Fig. 4A).

**Figure 4.**
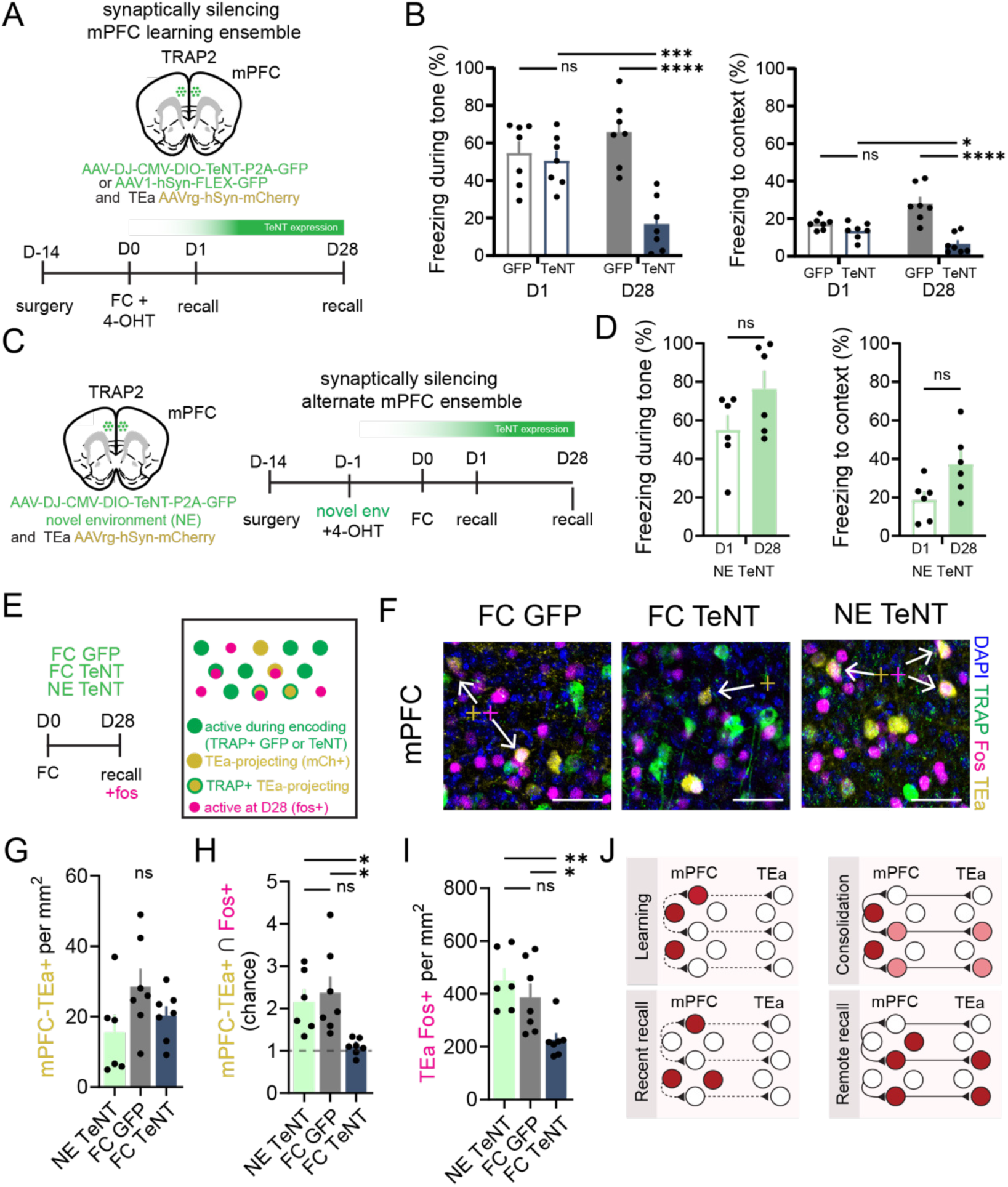
– Synaptic activity of mPFC learning-activated cells recruits mPFC-TEa neurons and is required for mote memory retrieval. **A**. Schematic of viral and experimental strategy to synaptically silence learning-activated PFC neurons. **B.** Left: tone freezing on D1 and D28 for all experimental groups (virus F(1,12)=2.5, P=0.13, Time 1,12)=35.9, P<0.0001, Virus x Time F(1,12)=25.7, P=0.0003, Two-way ANOVA with Sidak’s multiple comparisons st, n=7 GFP mice, n=7 TeNT mice). Right: context freezing on D1 and D28 for all experimental groups (virus 1,12)=0.5, P=0.4, time F(1,12)=38.6, P<0.0001, virus x time F(1,12)=18.8, P=0.001, Two-way ANOVA with Sidak’s ultiple comparisons test; n=7 GFP mice, n=7 TeNT mice). **C.** Same as A for synaptically silencing Novel Environment E)-activated mPFC neurons. **D.** Left: Tone freezing on D1 and D28 (paired t-test =0.15, n=6 mice/timepoint). Right: ntext freezing on D1 and D28 (paired t-test, P=0.12, n=6 mice/timepoint). Schematic of labeling strategy to examine the remote memory ensemble. **F.** Representative confocal images of mPFC across conditions. Arrows pointing toward double or triple positive cells, with color coded plus signs indicating expression. Scale bar = 50 µm. **G.** Density of TEa-projecting (mCh+) neurons for all three groups (one-way ANOVA 2,17)=2.2, P=0.13, n=6 NovEnv mice, n=7 GFP mice, n=7 TeNT mice). **H.** mPFC-TEa projecting neurons that were tivated during remote recall (Fos+) for all three groups (F(2,17)=5.9, P=0.01, one-way ANOVA with Tukey’s multiple mparisons test, n=6 NovEnv mice, n=7 GFP mice, n=7 TeNT mice). **I.** TEa neurons active during remote memory trieval (Fos+) (ANOVA F(2,17)=7.3, P=0.005, Tukey’s multiple comparisons tests, n=6 NovEnv mice, n=7 GFP mice, 7 TeNT mice). **J.** Cartoon summary of main findings. Color indicates neuronal involvement. Thickness of lines indicates strength of connection between neurons. <0.05, **P<0.01, ****P<0.0001. Error bars represent mean ± s.e.m.

Both TeNT-expressing mice (FC-TeNT) and GFP-expressing control mice (FC-GFP) exhibited high levels of freezing behavior during the recent memory test—conducted before the synaptic silencing likely took effect—indicating that both groups formed a robust memory. Strikingly, synaptically silencing mPFC learning-activated neurons during consolidation led to a strong reduction in freezing during the remote memory test (Fig. 4B). This suggests that ongoing synaptic transmission from learning-activated mPFC neurons is required to consolidate and express a remote memory.

To test the specificity of these findings, we used TRAP2 mice to drive TeNT expression in mPFC neurons activated by a different experience—exploration of a novel environment (NE)_—_and then performed FC in those mice the next day (Fig. 4C). In this case, both NE-TeNT and FC-GFP control mice had high levels of freezing during both recent and remote memory tests (Fig. 4D), indicating that synaptic activity of mPFC neurons specifically activated during fear learning is critical for remote memory formation. Because TeNT blocks synaptic transmission both during consolidation period and at the time of remote memory retrieval, we next investigated whether acute inhibition of learning-activated mPFC neurons would similarly impair remote memory retrieval. To test this, we expressed an inhibitory opsin in learning-activated mPFC neurons, and silenced them during memory retrieval (Extended Data Fig. 7A). Opsin-expressing mice showed a small but significant decrease in cued freezing during laser-on trials (Extended Data Fig. 7B), although they still froze significantly more than FC-TeNT mice (Extended Data Fig. 7C). Taken together, these results demonstrate that synaptic activity in mPFC neurons activated during learning is required during the consolidation period to establish a lasting memory trace.

Next, we investigated whether synaptic silencing of mPFC neurons active during learning impairs the recruitment of mPFC-TEa neurons into the remote memory trace (Fig. 4E,F). First, we confirmed that we labeled similar numbers mPFC-TEa neurons (mCherry+) neurons between groups (Fig. 4G). We then used immunofluorescence to quantify the number of active (Fos+) mPFC-TEa neurons during remote memory retrieval, comparing FC-TeNT to NE-TeNT and FC-GFP control groups. FC-TeNT animals had fewer Fos+ mPFC-TEa neurons and fewer Fos+ cells in TEa itself (Fig. 4H,I). Additionally, FC-TeNT-expressing mice had fewer Fos+ mPFC neurons during remote retrieval compared to NE-TeNT and FC-GFP controls, despite having a similar number of TRAP+ cells (Extended Data Fig. 8B,C). These data indicate that blocking synaptic communication from learning-activated mPFC neurons disrupts the recruitment of mPFC-TEa neurons and reduces overall mPFC and TEa activation during remote memory retrieval.

To gain further insight into the time-dependent recruitment of mPFC-TEa neurons, we examined the composition of the remote recall-activated mPFC-TEa neurons with regards to their activation during learning. Specifically, we calculated the proportion of mCh+/TRAP+/Fos+ cells (i.e., neurons projecting to TEa that were activated during both learning and remote recall) relative to the total mCh+/Fos+ population (i.e., projecting to TEa and activated during remote recall). We termed this ratio the ‘Reactivation Index’, where higher numbers indicate a more static memory trace (i.e. the same cells were more likely to be activated during learning and remote memory). While there were no significant differences in the number of TRAP+ mPFC-TEa neurons (Extended Data Fig. 8D), FC-TeNT animals exhibited a significantly higher Reactivation Index compared to both control groups (Extended Data Fig. 8E). This finding suggests that synaptic inhibition of learning-activated mPFC neurons prevents the recruitment of new mPFC-TEa neurons and results in a more static memory trace. Overall, these results indicate that ongoing synaptic activity of learning-activated mPFC neurons is necessary during memory consolidation to recruit mPFC-TEa neurons—and TEa itself—to support remote memory recall (Fig. 4J).

## Discussion

While long-standing theories of memory consolidation have proposed that remote memories are stored in a distributed cortical network^10^, it has remained unclear how these networks are constructed. Here we identify a specific class of mPFC projection neurons—mPFC-TEa neurons—that are gradually recruited to a memory trace over time. These neurons increasingly encode a conditioned cue and conditioned freezing and develop population dynamics that distinctly represent the intersections of conditioned cues, contexts, and fearful behaviors. The mPFC-TEa pathway is selectively required for remote memory retrieval, whereas a subcortical projection (mPFC-BLA) is necessary only for recent memory retrieval. We further show that ongoing synaptic activity from learning-activated mPFC neurons is essential for recruiting mPFC-TEa neurons and for successful consolidation of a remote memory. Together, our findings demonstrate that memory encoding initiates synaptic processes in the mPFC that gradually recruits a distinct class of cortico-cortical projection neurons into the memory trace. We thus provide a mechanistic framework for the formation of a distributed cortical network during systems consolidation.

Previous studies suggests that mPFC neurons that are active during learning enter a quiescent period during early consolidation, followed by a functional maturation that enables them to support remote memory retrieval^13,14^. This maturation process involves growth of dendritic spines—sites of synaptic contact— and strengthened synaptic connections with other mPFC neurons that were active during learning^13,14^. Studies have also shown that new cortical neurons are gradually incorporated into the memory trace^13–15^, although their identity has remained unclear. Our findings using longitudinal Ca²⁺ imaging reveal that mPFC-TEa neurons with strong shock-evoked responses during learning became largely nonresponsive or suppressed during tone presentations at remote memory retrieval, whereas neurons suppressed during foot shocks were more likely to become tone-responsive later. In complementary experiments, we demonstrate that many mPFC-TEa neurons activated (Fos+) during remote memory were not previously activated (TRAP-) during learning, and that synaptic inhibition of learning-activated mPFC neurons prevented this time-dependent recruitment. These findings indicate that functional recruitment of mPFC-TEa neurons is critical to establish a lasting memory trace. Thus, our study uncovers additional cell type-specific mechanisms of memory consolidation, extending beyond the persistent roles of neurons excited during learning.

Cortical association areas such as TEa have long been implicated in remote memory ^21–23,37–41^, yet the mechanisms by which TEa is recruited into the remote memory trace have remained unclear. TEa encodes conditioned stimuli and emotional behavioral responses^42^ and contributes to remote auditory fear memory through its projections to the amygdala^22,23,37,38^. Our findings build on this by revealing a time– and learning-dependent increase in both mPFC-TEa activity and in TEa activity during conditioned tone presentations. Compared to the broader mPFC population, mPFC-TEa neurons were enriched in functional clusters that shifted their neural dynamics from recent to remote recall. Although these neurons represent only a subset of the population, they played a key role in shaping overall network dynamics. *In silico* lesioning of these cells abolished the distinct population-level representation of tone-freezing epochs at the remote timepoint. Theoretical attractor models of memory propose that such orthogonalization reduces memory interference and enhances memory robustness^43–46^. Thus, mPFC-TEa dynamics likely shape TEa activity during remote recall and may serve to stabilize memory representations across the neocortex.

TEa is a defining projection target of mPFC IT neurons^19^. Consistent with this, our whole-brain axon tracing revealed that mPFC-TEa neurons send dense axon collaterals to classic IT targets including the RSP, INS, AUDv, PTLp, BLA, CLA and NAc. Notably, mPFC-TEa neurons activated during memory retrieval preferentially targeted the claustrum—an understudied region that is similar to neocortex in developmental origin but lacks canonical cortical architecture^47^. The claustrum’s broad cortical connectivity has been implicated in fear memory consolidation^48^. While our study focused on the mPFC-TEa pathway, RSP^49^, INS^50^, AUDv^51^, PTLp^52^ also have established roles in systems consolidation and remote memory retrieval, so these other mPFC pathways may likewise have dynamic roles in memory retrieval. Top-down mPFC control of sensory areas could support attentional control of cortical processing, whereas projections to the striatum and amygdala may more directly regulate behavioral outputs. Our results suggest a model in which local synaptic interactions recruit mPFC IT neurons into a memory trace, serving as a key mechanism by which the brain constructs distributed cortical networks to support long-term memory storage.

Overall, our findings reveal a dynamic mechanism of memory consolidation in the mPFC that integrates synaptic, cellular, and circuit-level changes to support the formation of a lasting memory. These insights are significant because in clinical contexts, remote fear memories of traumatic events are often more resistant to therapeutic intervention than recent memories^53–55^. A detailed understanding of the circuits and cell types involved in memory consolidation and retrieval—and how they evolve over time—may uncover novel therapeutic targets and lay the groundwork for more effective treatments.

## Methods

All experiments were conducted in accordance with procedures established by administrative panels on laboratory animal care at the University of California, Los Angeles.

## Animals

Female and male C57B16/J mice (JAX Stock No. 000664) or TRAP2 mice (JAX Stock No. 030323) were group housed (2–5 per cage) and kept on a 12-hour light cycle (lights on 7am-7pm). Sex was determined by visual examination of external genitalia at weaning. All animal procedures followed animal care guidelines approved by the University of California, Los Angeles Chancellor’s Animal Research Committee.

## Behavioral Assays

### Fear conditioning

Animals were habituated to the behavior arena and tones one day before fear conditioning (D-1). The tone (4000Hz) was presented for 10 seconds, four times. The inter-tone interval for all days was between 40 and 90 seconds. On the day of fear conditioning (D0), animals were placed in the same environment and exposed to 8 presentations of the 10 second tone (CS) which co-terminated with a one second, 0.55mA shock. For both recent (D1) and remote (D28) recall, animals were placed in a separate context and were exposed to four presentations of the CS for 30 seconds. For optogenetic experiments, a red laser (635nm, Laser Century) was on for the duration of interleaved tones. Overhead videos were recorded (Point Grey Chameleon3) for freezing analysis. Videos were processed using markerless point tracking^56^ which was then used to calculate freezing using pose estimation algorithms in BehaviorDEPOT^57^.

### Fiber photometry

Recordings were performed using a commercial fiber photometry system (RZ10x, Tucker Davis Technologies) with two excitation wavelengths, 465 and 405 nm, modulated at 211 and 566 Hz respectively. Light was filtered and combined by a fluorescent mini cube (Doric Lenses). Emission was collected through the mini cube and focused onto a femtowatt photoreceiver (Newport, Model 2151).

Samples were collected at 1017 Hz and demodulated by the RZ10x processor. Time stamps to synchronize experimental events and recordings were sent via TTLs to the RZ10x system via Arduino, controlled by custom MATLAB (MathWorks) code.

### 4-Hydroxytamoxifen Preparation and Delivery

4-hydroxytamoxifen (4-OHT; Sigma, Cat# H6278) was dissolved at 20 mg/mL in ethanol by shaking at 37°C for 15 min and was then aliquoted and stored at –20°C for up to several weeks. Before use, 4-OHT was redissolved in ethanol by shaking at 37°C for 15 min, a 1:4 mixture of castor oil:sunflower seed oil (Sigma, Cat # S259853 and S5007) was added to give a final concentration of 10 mg/mL 4-OHT, and the ethanol was evaporated by vacuum under centrifugation. The final 10 mg/mL 4-OHT solutions were always used on the day they were prepared. All injections were delivered intraperitoneally (i.p.) after fear conditioning (D0) or novel environment exploration (D-1) to deliver 50 mg/kg of 4-OHT to subjects.

After 4-OHT administration, animals were single-housed in a quiet room for at least six hours before being returned to their home cages.

### Miniscope recordings

Mice were handled and habituated to the microscope for 5 days before the start of the experiment. During recording sessions, a miniscope (V4.3) was secured to the baseplate and the mice were allowed to acclimate in their home cage for 5 min. Miniscope imaging took place throughout the entire session.

Behavior was simultaneously recorded using miniscope recording software to synchronize the data streams

## Surgery

### Virus injection and implant surgeries

All surgical procedures used isoflurane anesthesia and a stereotaxic frame (Kopf 963). Heads were sterilized with alternating scrubs of betadine and 70% ethanol. Eye moisture was maintained with ophthalmic ointment. For pain management, mice received 5 mg/kg carprofen diluted in 0.9% saline subcutaneously. Mice received one carprofen injection during surgery and daily injections for 2 days following surgery. For surgeries with injections and no implants, following injections, skin was sutured. For implant surgeries, all implants were secured to the skull with C&B Metabond (Parkell) with any exposed skull covered as well.

For optogenetic experiments, AAV5-hSyn-Jaws-KGC-GFP-ER2 (10e12 GC/mL, 500nL at 100nL/min, Addgene #65014) was injected bilaterally into PL subregion of mPFC (referred to as mPFC) (+2 AP, 0.35 ML, –2 DV mm from bregma). Fibers (100 um diameter, Newdoon) were implanted over BLA (–1.1 AP, 3.1 ML, 4.9 DV mm from bregma) or TEa (–2.9 AP, 4.4 ML, 3.3 DV mm from bregma).

For fiber photometry experiments, AAV5-syn-jGCaMP8m-WPRE (10e12 GC/mL, 500nL at 100nL/min, Addgene #162375) was injected into TEa with a fiber (200um diameter, Newdoon) implanted 200um above the injection site.

Miniscope experiments were performed as previously described^58,59^. Briefly, AAV8-syn-jGCaMP7f-WPRE or AAV8-syn-FLEX-jGCaMP7f-WPRE (10e13 GC/mL, 500nL at 100nL/min, Addgene #104488) was injected into the left PL region of mPFC. For mPFC-TEa experiments, AAVrg-Ef1a-mCherry-IRES-Cre was injected into the left TEa. A week later, a 1mm craniotomy was made around the mPFC injection site and the cortex was aspirated until 200um above the injection site with a constant flush of saline. A 1mm GRIN lens (Inscopix) was implanted into the craniotomy and secured. The GRIN lens was covered with Kwik-Sil (World Precision Instruments). Animals received dexamethasone (0.2 mg/kg) for 5 days following surgery.

For synaptic inhibition experiments, tetanus toxin light-chain (AAV-DJ-CMV-DIO-EGFP-2A-TENT, 10e12 GC/mL, 500nL at 100nL/min) (Stanford Gene Vector and Virus Core), or GFP (AAV1-hSyn-DIO-EGFP, 10e12 GC/mL, 500nL at 100nL/min) (Addgene #50457), or stGtACR2 (AAV1-hSyn1-SIO-stGtACR2-FusionRed, 500nL at 100nL/min) (Addgene #105677) was injected in mPFC and AAVrg-Ef1a-mCherry-IRES-Flpo (10e12 GC/mL, 300nL, 50nL/min, Addgene #55634) was injected into TEa using TRAP2 mice.

For stGtACR2 experiments, fibers were placed bilaterally 100um dorsal to the injection site.

For brain clearing and imaging experiments of TRAP2 animals, AAV8-hSyn-Con/Fon-hChR2(H1234R)-EYFP (10e13 GC/mL, 500nL, 100nL/min, Addgene #55645) was injected into the left mPFC and AAVrg-Ef1a-mCherry-IRES-Flp was injected into the left TEa. For wildtype animals, AAV8-EF1a-double floxed-hChR2(H134R)-EYFP-WPRE-HGHpA (10e13 GC/mL, 500nL, 100nL/min, Addgene #20298) was injected into the left mPFC and AAVrg-Ef1a-mCherry-IRES-Cre (10e12 GC/mL, 300nL, 50nL/min, Addgene #55632) was injected into the left TEa.

### Brain slice histology and immunostaining

Mice were deeply anesthetized with isofluorane and perfused intracardially with 20mL of PBS followed by 4% paraformaldehyde (PFA, Electron Microscopy Sciences) on ice. Brains samples were removed and fixed in 4% PFA overnight, and were then moved to PBS. Brains were then sectioned via vibratome at 60 microns to perform immunohistochemistry. To detect Jaws expression, anti-GFP was used (1:2000, AVES Labs, GFP 1020). To detect mCherry, rat anti-mCherry (1:1000, M11217, Rockland) was used.

To detect cFos, anti-cFos was used (1:1000, Synaptic Systems 226 003). Sections were incubated in primary antibody overnight at 4°C, except cFos primary, which incubated for 48 hours. Sections were then rinsed and incubated in corresponding secondary antibody (1:2000) at room temperature for 2 hours. Sections were washed and stained with DAPI before being mounted. Images of fluorescent expression and implant targeting were taken using Leica DM6 scanning microscope (Leica Microsystems) using a 10x objective and a confocal microscope (Leica). For each animal, 3-4 sections were counted and then averaged across sections. Overlap of Fos+ and mCherry+ was calculated relative to chance levels, defined as (Fos+mCh+/DAPI) /(Fos+/DAPI)x(mCh+/DAPI)).

### Miniscope data processing and analysis

Cell footprints and Ca^2+^ fluorescence time series were extracted from miniscope recordings using Minian^60^ with manual validation of cell footprints and signals. Videos were recorded at 20Hz and analyzed at full spatial and temporal resolution. Accepted neurons and their calcium activity traces were exported to MATLAB for further analysis.

To identify neurons that encoded tone or freezing information, we evaluated single cell responses during tone presentation and freezing behavior. Using the MATLAB *perfcurve* function, we generated receiver operating characteristic curves (ROC) for individual neurons and measured the area under the curve (auROC). To determine if a neuron encoded an event, we generated a null distribution of auROCs by circularly shuffling event timing and recalculating the auROC over 1000 permutations. Neurons were considered significantly excited by an event if their auROC was greater than 97.5% of auROCs in the null distributions and significantly suppressed if their auROC value was in the lowest 2.5% of all auROCs in the null distribution. To account for differences in total number of recorded neurons, modulated neurons are reported as a percentage of excited or suppressed neurons relative to the total number of recorded neurons. Average number of neurons recorded in mPFC-TEa recordings was 17.03 +/-0.03 SEM. Average number of neurons recorded in mPFC recordings was 77.8 +/-1.9 SEM.

### Tracking

Tracking mPFC-TEa neurons across recording sessions was done using CellReg^61^ followed by manual validation of alignment. One subject was excluded from alignment analyses due to optical misalignment across recording days. A vector describing shock responsiveness of each cell and a corresponding vector describing tone responsiveness of each cell were created, calculating the observed overlap. A permutation test was used to determine the significance of shock and tone response overlap, randomly shuffling the tone responsiveness vector 1000 times, re-computing overlap for each permutation to calculate a null distribution. The observed overlap was compared against the null distribution to obtain a p-value, achieving significance by being in the top or bottom 2.5% of permuted null distribution values.

### Clustering

Calcium imaging data were standardized using *sklearn.preprocessing.StandardScaler* before calculating the average tone response for each recording. All recordings from both mPFC and mPFC-TEa groups were combined into Day 1 or Day 28 sessions. Pairwise distances between averaged tone responses were computed using Pearson’s correlation distance (*scipy.spatial.distance.pdist*). The optimal number of clusters was identified using the KneeLocator method (*kneed.KneeLocator*). Hierarchical clustering was performed with Ward’s linkage (*scipy.cluster.hierarchy.linkage*). A permutation test was used to determine the significance of observed clustering effects. Specifically, group labels (mPFC or mPFC-TEa) of each leaf were randomly shuffled 1000 times, and clustering was re-computed for each permutation to calculate a null distribution. The observed clustering metric was compared against the permuted distribution to obtain a p-value, achieving significance by being in the top or bottom 2.5% of permuted null distribution values.

### Embeddings

Behavior-informed embeddings were generated with CEBRA^31^. Models were fit using calcium signals with tone and/or freezing vectors, which were then used to plot latent activity. To generate the models, batch_size, learning_rate, temperature, max_iteration, distance, and time_offsets parameters were consistent between models. Due to variance in the number of neurons recorded between animals, output dimension parameter was selected for each recording by generating a scree plot and finding the elbow using the Python packages *scipy* and *sklearn*. InfoNCE loss plots, a measure of goodness of model fit^31^, were generated using *cebra.plot_loss* function. Distance between centroids was calculated by taking the mean of each class and calculating the distances.

For *in silico* lesions, embeddings were regenerated and centroid distances recalculated after removing neurons belonging to the lesioned cluster. For shuffled control lesion, an equivalent number of neurons for each recording that belonged to clusters other than the previously lesioned one were randomly selected and removed.

## Brain-clearing and whole-brain imaging

Mouse brain tissue was prepared following a modified version of the Adipo-Clear protocol^19,62^. Briefly, mice were intracardially perfused on ice with 20 mL phosphate-buffered saline (PBS; Invitrogen) followed by 4% paraformaldehyde (PFA; Electron Microscopy Sciences). Brains were then hemisected approximately 1 mm lateral to the midline and post-fixed overnight at 4°C in 4% PFA. The following day, samples were sequentially dehydrated using methanol (MeOH, Fisher Scientific) mixed with B1n buffer (1:1000 Triton X-100, 2% w/v glycine, 1:10,000 NaOH 10N, 0.02% sodium azide). Each methanol gradient (20%, 40%, 60%, and 80%) was applied for 1 hour using a nutator (VWR). Samples were then rinsed twice with 100% MeOH for 1 hour each and subsequently incubated overnight in a 2:1 dichloromethane (DCM solution. The next day, samples were washed twice in 100% DCM for 1 hour each, followed by three washes in 100% MeOH for progressively longer durations (30 minutes, 45 minutes, and 1 hour). Tissue samples were then bleached for 4 hours in a 5:1 H₂O₂ solution. Rehydration was achieved through a series of MeOH/B1n buffer washes in decreasing methanol concentrations (80%, 60%, 40%, and 20%) for 30 minutes each, followed by a final 1-hour wash in B1n buffer.

Permeabilization was carried out with 5% DMSO/0.3 M glycine in PTxWH buffer for 1 hour, followed by an additional 2-hour incubation in fresh permeabilization solution. Samples were then rinsed in PTxwH for 30 minutes and left in fresh PTxwH buffer overnight. On the following day, two more PTxwH washes (1 hour and 2 hours) were performed.

Samples were incubated with primary anti-GFP antibody (AVES Labs, GFP 1020) at a dilution of 1:2000 in PTxwH, with continuous shaking at 37°C for 11 days. This was followed by sequential PTxwH washes: twice for 1 hour each and twice for 2 hours each, with additional PTxwH exchanges over a 2-day period at 37°C. Following primary antibody incubation, samples were incubated with secondary antibody at 1:1500 (AlexaFluor 647, ThermoFisher Scientific) at 37°C for 8 days, with regular PTxwH washes over two days. After antibody staining, samples were rinsed in PBS twice (1 hour each), then for 2 hours twice, and left overnight. Dehydration involved a graded series of MeOH washes (20%, 40%, 60%, and 80%) for 30 minutes each, followed by three 100% MeOH washes (30 minutes, 1 hour, and 1.5 hours).

Samples were incubated overnight in a 2:1 DCM solution on a nutator. The next day, samples were washed twice in 100% DCM for 1 hour each, then cleared in 100% dibenzyl ether (DBE), with DBE refreshed after 4 hours. Cleared samples were stored in DBE at room temperature in darkness, and imaging was performed after at least 24 hours.

### Whole-brain imaging

Brain samples were imaged using a light-sheet microscope (Ultramicroscope II, LaVision Biotec) outfitted with a sCMOS camera (Andor Neo) and a 2x/0.5 NA objective lens (MVPLAPO 2x) with a 6-mm working distance dipping cap. Image stacks were captured at 0.8x optical zoom and controlled through Imspector Microscope v285 software. For axonal imaging, 488-nm (20% laser power) and 640-nm (50% laser power) lasers were used. Scanning was performed with a 3 µm step size, employing a continuous light-sheet scanning method with a contrast-adaptive algorithm for the 640-nm channel (20 acquisitions per plane), and without horizontal scanning for the 488-nm channel.

### Whole-brain analysis

DeepTRACE^19^ was used to align and quantify projections across the brain. Briefly, regional axon innervation was quantified using MATLAB (MathWorks) by calculating the density of skeletonized pixels within each brain region. Pixels exceeding a threshold of 64 were counted, then normalized by the total pixel count of the respective region. To account for variations in total fluorescence intensity and viral expression, this regional pixel count was further normalized by the total number of labeled pixels across the brain. Regions were defined based on a modified version of the LSFM atlas. The atlas was cropped at the anterior and posterior boundaries to align with the visible tissue extent in our data, and analyses excluded fiber tracts, ventricular systems, cerebellum, and olfactory bulb. Regional axon labeling comparisons between groups are presented as unpaired t-tests with no corrections for multiple comparisons. For each group, the axon counts were averaged, and the mean values with SEM were plotted.

Statistical analyses were conducted with two-tails and performed in Python, MATLAB, and GraphPad Prism v9.

## Code availability

Custom MATLAB requests for analyzing data are available upon request.

## Data availability

Data are available in the main text and extended data materials.

## Acknowledgements

We thank Drs. Dean Buonomano, David Glanzman, Emilie Marcus, Megha Segal, Zachary Pennington and Andre de Sousa for helpful comments on the manuscript. ZZ was supported by F32MH133387. AY was supported by an ARCS Foundation Fellowship. This work was supported by R01MH137461 and a Vallee Scholars Award to LAD.

## Author Contributions

ZZ performed all experiments and analyzed the data except for the PL-BLA optogenetics experiments. AY performed the PL-BLA optogenetics experiments. MPS and RH assisted with experiments. LAD and ZZ supervised the project. LAD, ZZ and AY acquired funding. LAD and ZZ wrote the original draft. LAD, ZZ and AY edited the manuscript.

## Competing Interests

Authors declare that they have no competing interests.

**Extended Data Figure 1.**
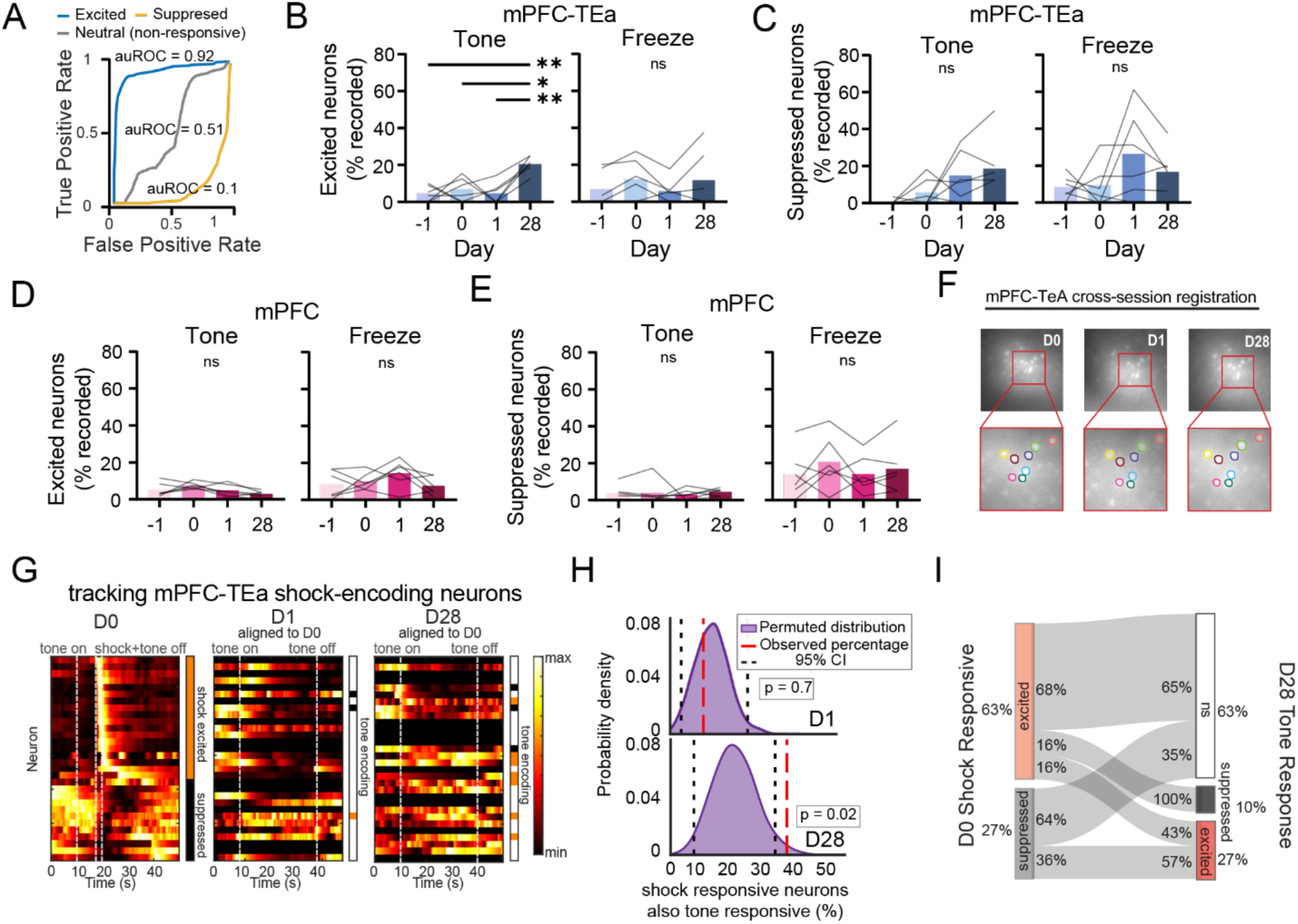
– Detailed analyses of tone, freeze, and shock modulated neurons. **A**. Example Receiver Operating Characteristic (ROC) curves for neurons that are excited (blue), suppressed (yellow), or non-responsive (gray) to the tone. Quantification of the area under the ROC curve (auROC) overlaid on the curve. **B.** Proportion of mPFC-TEa neurons excited by tone or freezing across time (Tone excited: F(2.2, 11.2)=10.3, P=0.002, Repeated Measures ANOVA with Tukey’s multiple comparisons tests. Freeze excited: F(1.7, 8.7)=1.4, P=0.29, Repeated Measures ANOVA, n=6 mice). **C.** Same as B for suppressed neurons (Tone suppressed: F(1.3, 6.9)=3.5, P=0.09, Repeated Measures ANOVA; Freeze suppressed: F(1.7, 8.9)=2.2, P=0.16, Repeated Measures ANOVA). **D.** Same as B for mPFC neurons (Tone excited: F(1.9, 9.7)=2.4, P=0.13, Repeated Measures ANOVA, Freeze excited: F(1.5,7.9)=1.2, P=0.31, Repeated Measures ANOVA, n=6 mice). **E.** Same as C for mPFC neurons (Tone suppressed: F(1.2, 6.3)=0.1, P=0.8, Repeated Measures ANOVA; Freeze suppressed: F(1.5, 7.9)=1.1, P=0.35, Repeated Measure ANOVA, n=6 mice). **F.** Example images of registering the same neurons across sessions in mPFC-TEa recordings. **G.** Heatmap showing tone-averaged response of shock-encoding neurons (left), aligned to responses at recent recall (middle), and remote recall (right). Orange bar indicates cells excited by shock or tone, black bar indicates cells suppressed by shock or tone. **H.** Overlap between shock-encoding during learning (D0) and tone-encoding during recent recall occurs (13%) and remote recall (33%). Permuted distribution of randomized overlap between shock– and tone-encoding in purple, observed proportion of overlap in red. 95% confidence interval in black. **I.** Sankey plot visualizing outcome of mPFC-TEa shock-encoding neurons at remote recall. *P<0.05, **P<0.01.

**Extended Data Figure 2.**
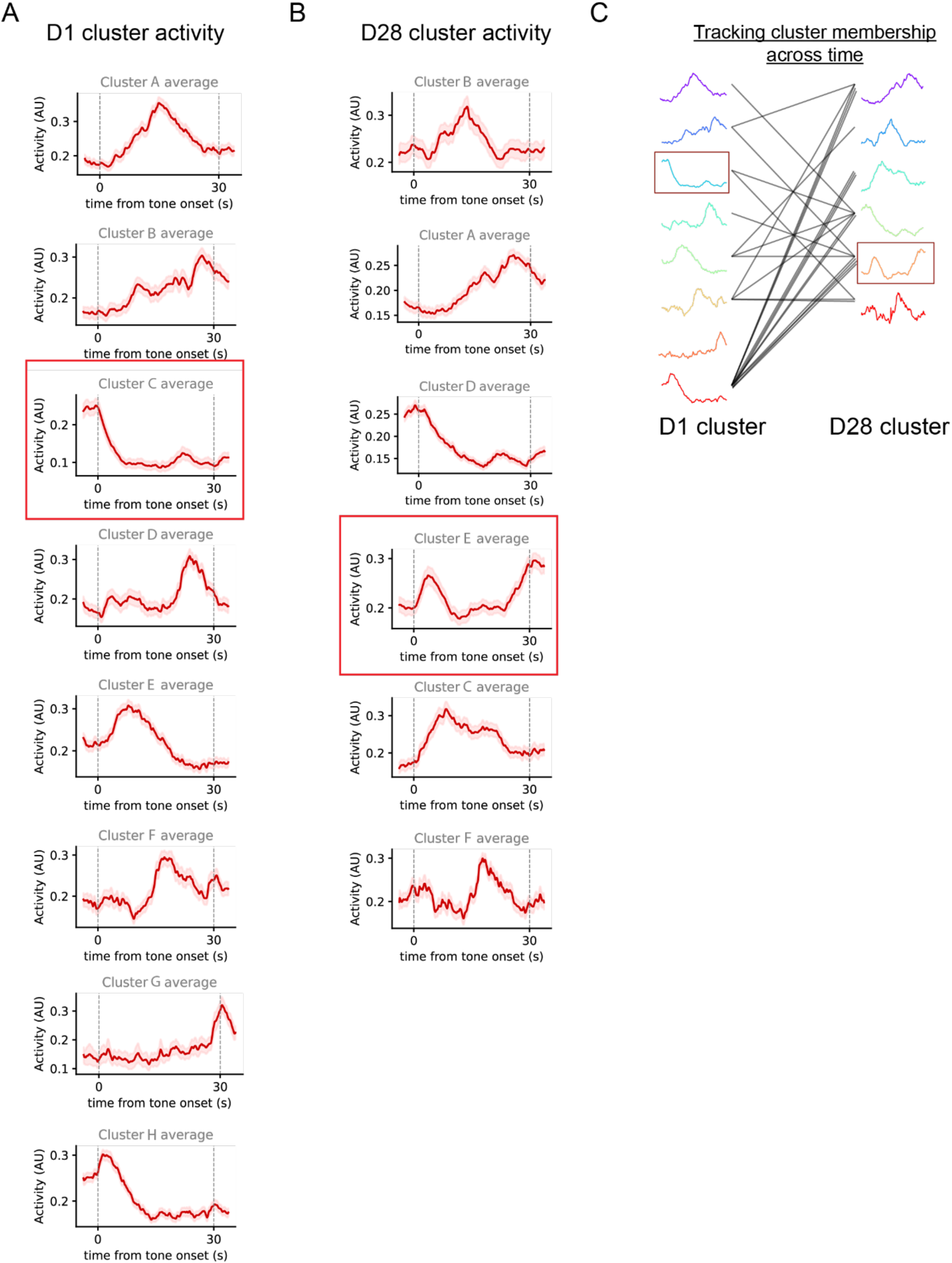
– Averaged tone response of functional clusters. **A**. Average response of hierarchically grouped mPFC and mPFC-TEa neurons clusters during recent recall. Red outlined cluster indicates higher than expected mPFC-TEa membership. **B.** Same as A for tone response during remote memory. Cluster IDs correspond to labels in Figure 2, but are organized to qualitatively match a cluster from A.. C.. Cluster membership of mPFC-TEa neurons aligned between D1 and D28 recall. n=25 neurons.

**Extended Data Figure 3.**
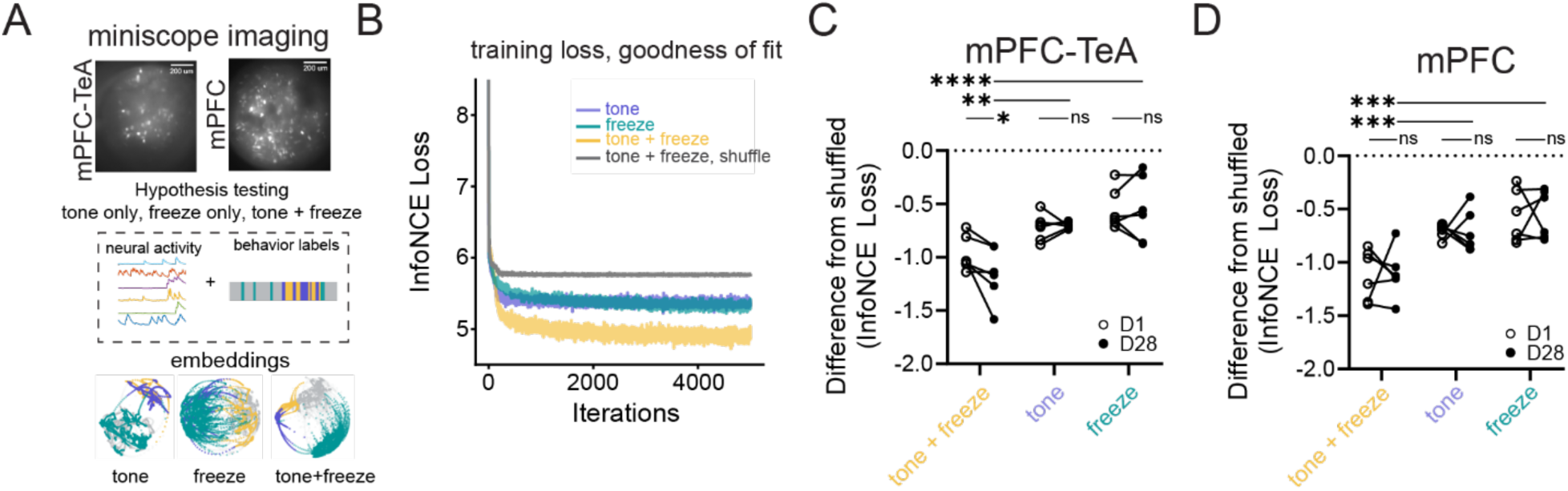
– Tone and freezing activity capture mPFC-TEa and mPFC population activity. **A**. Summary of population activity analyses. Top: example field of views from miniscope imaging in mPFC-TEa neurons or general mPFC neurons. Middle: example activity traces and example behavioral labels. Bottom: example embeddings trained on tone, freezing, or tone and freezing information. B. Representative InfoNCE loss across iterations of training for embeddings with tone (purple), freeze (green), tone and freeze (yellow), or shuffled tone and freeze (gray) variables. InfoNCE loss is a metric of embedding quality with lower scores indicating more efficient embeddings. C. For mPFC-TEa recordings, InfoNCE loss for embeddings trained on combined tone and freezing, tone only, or only freezing information (Behavior F(2,15)=12.5, P=0.0006; Time F(1,15)=0.9, P=0.34; Behavior x Time F(2,15)=4.7, P=0.026, two-way ANOVA with Sidak’s multiple comparisons test, n=6 mice). D. Same as C for mPFC recordings (Behavior F(2,15)=22.5, P<0.0001; Time F(1,15)=0.005, P=0.9; Behavior x Time F(2,15)=0.01, P=0.9, two-way ANOVA with Sidak’s multiple comparisons test, n=6 mice). * P<0.05, **P<0.01, ***P<0.001, ****P<0.0001

**Extended Data Figure 4.**
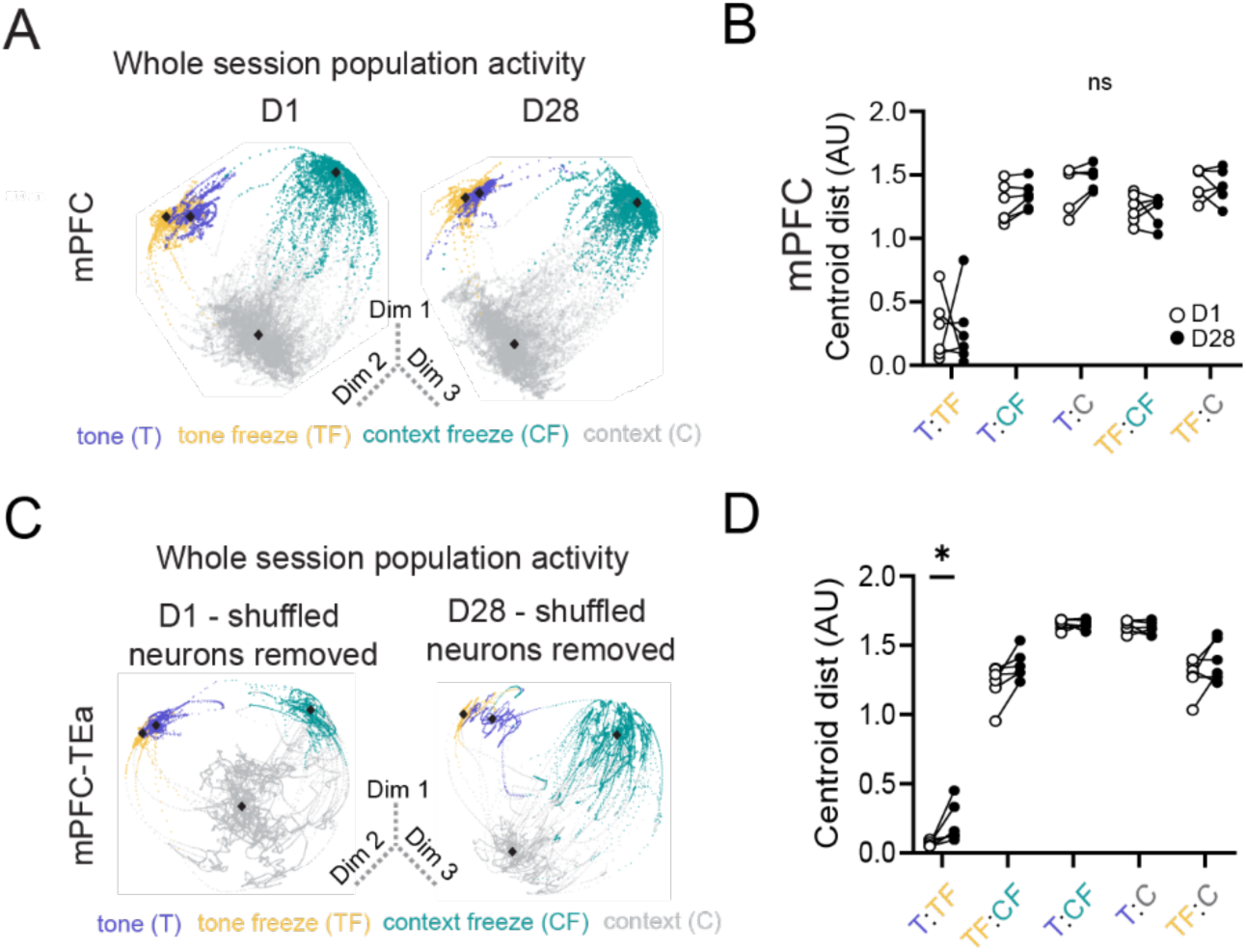
– Additional population analyses for mPFC-TEa and mPFC neurons. **A**. Example embeddings of datafrom mPFC recordings with their full dataset. **B.** Distance between centroids at D1 vs. D28 for mPFC embeddings (Centroids F(4,25)=97.8, P<0.0001, Time F(1,25)=0.2, P=0.6, Centroids x Time F(4,25)=0.2, P=0.9, Two-way Repeated Measures ANOVA, n=6 mice). C. Example embeddings of data from an mPFC-TEa recordings in which the Cluster lesions were shuffled for both Day 1 and D28. D. Distance between centroids at D1 vs. D28 for mPFC-TEa shuffled lesion embeddings (Centroids F(4,25)=244.1, P<0.0001, Time F(1,25)=5.9, P=0.02, Centroids x Time F(4,25)=2.8, P=0.03, Two-way Repeated Measures ANOVA, Šídák’s multiple comparisons test, n=6 mice). *P<0.05.

**Extended Data Figure 5.**
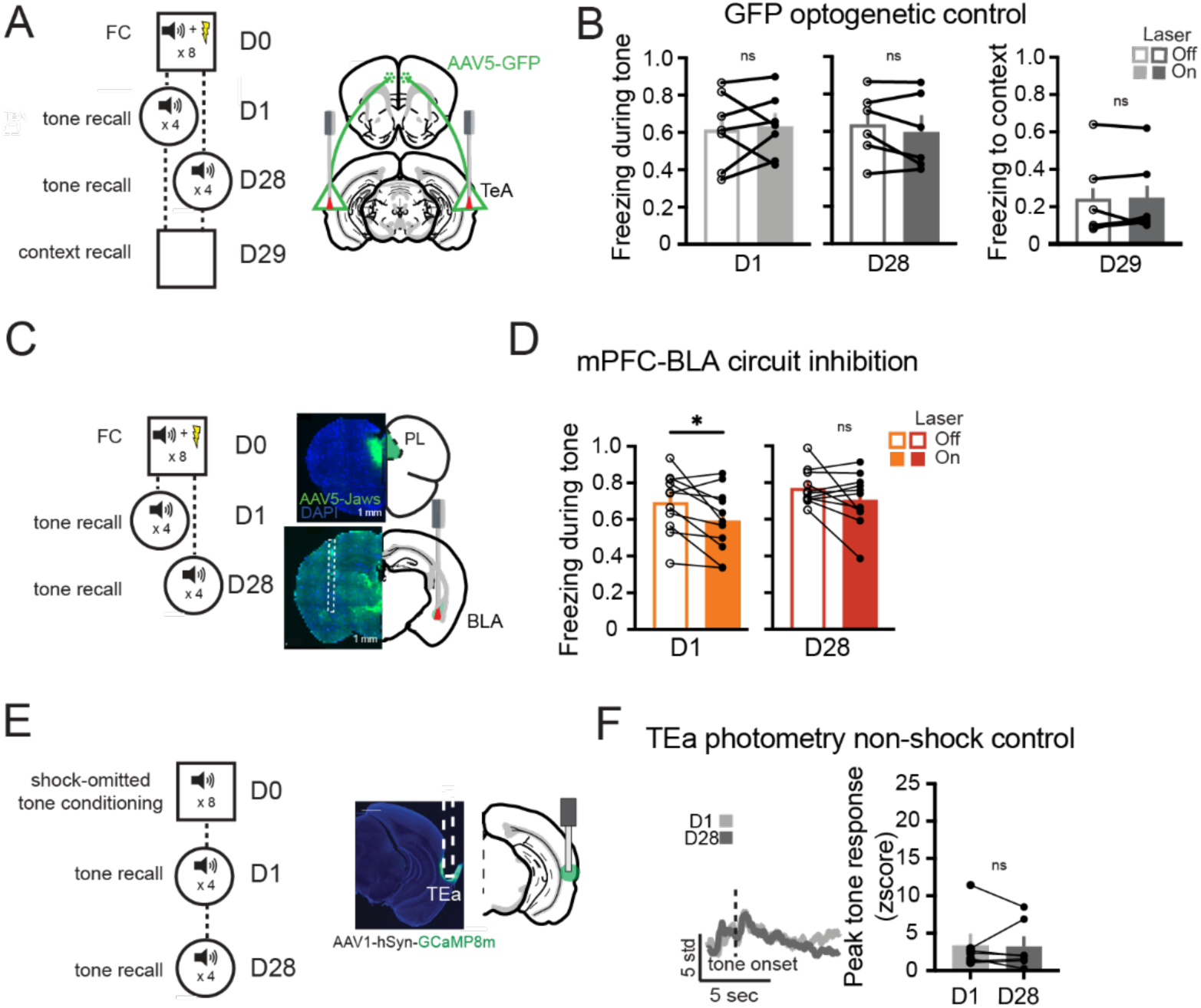
– Control experiments for optogenetic manipulations and TEa fiber photometry. **A**. Schematic of strategy to test fluorophore control virus in mPFC-TEa circuit manipulation during memory recall. B.. Left: D1 tone freezing in GFP control mice (P=0.7, paired t-test, n=6 mice). Middle: D28 tone freezing (P=0.26, paired t-test, n=6 mice). Right: context freezing on D29 (P=0.7, paired t-test, n=6 mice). C. Schematic of strategy to inhibit mPFC axons in BLA during D1 and D28 memory retrieval. D. Tone-induced freezing at D1 and D28 recall with and without laser (D1: P=0.15, D28: P=0.02, paired t-tests, n=11 mice each). **E**. Schematic of experimental and surgical strategy to test TEa response to tones presented without shock. F. Left: Representative traces of recent and remote TEa responses to tone onset. Right: tone-evoked TEa activity in non-shocked control mice on D1 and D28 (P=0.8, paired t-test, n=7 mice). *P<0.05. Error bars represent mean ± s.e.m.

**Extended Data Figure 6.**
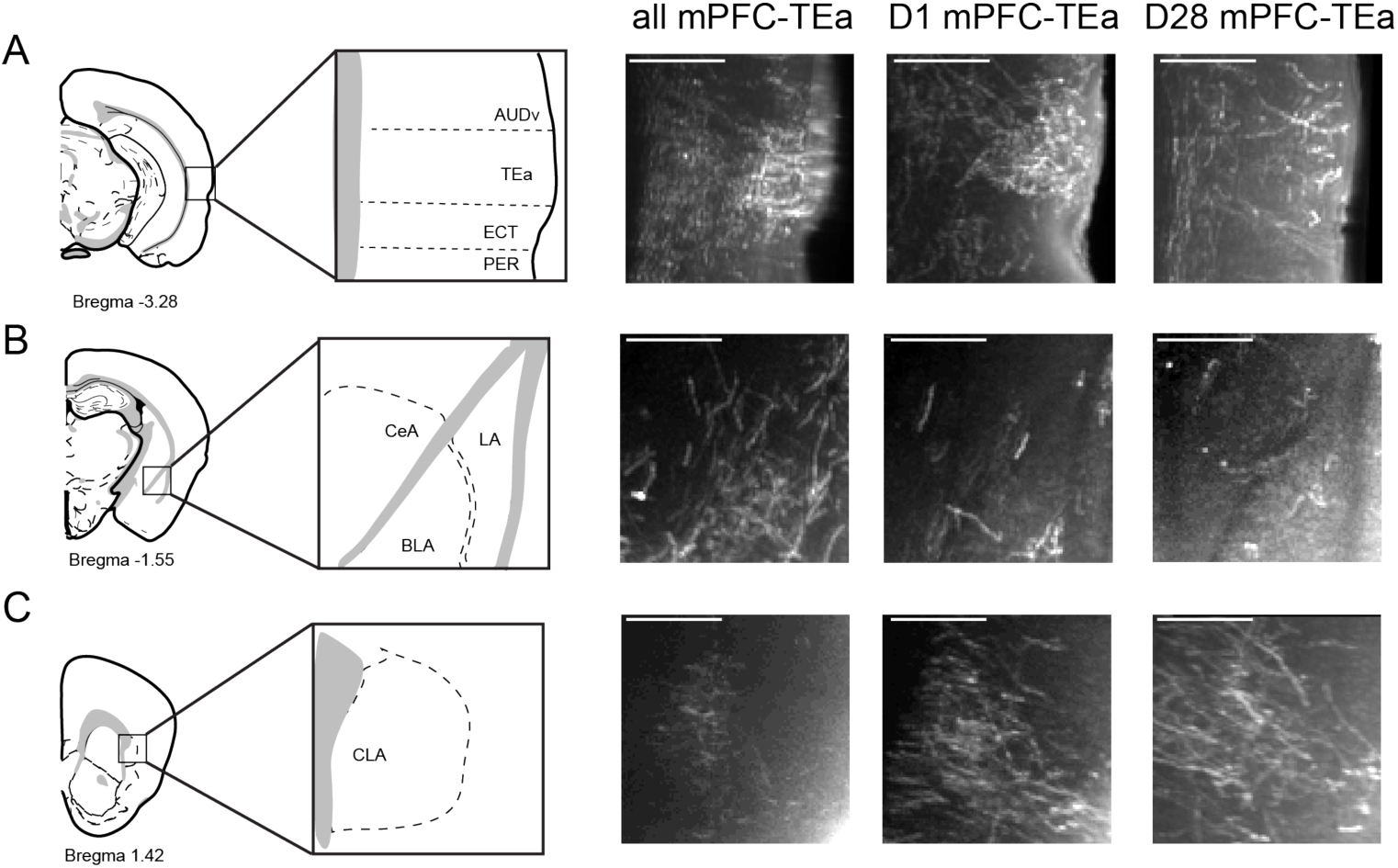
– Raw images of whole brain axon labeling. **A**. (left) Atlas schematic of coronal section with zoom-in showing location of TEa. (right) Representative images from mPFC-TEa, D1 TRAP, D28 TRAP groups. Images are max projections (30 um). **B**. Same as A for amygdala. C. Same as A for claustrum. AUDv, ventral auditory cortex. TEa, temporal association cortex. ECT, ectorhinal cortex. PER, perirhinal cortex. LA, lateral amygdala. BLA, basolateral amygdala. CeA, central amygdala. CLA, claustrum. Scale bars, 500 um.

**Extended Data Figure 7.**
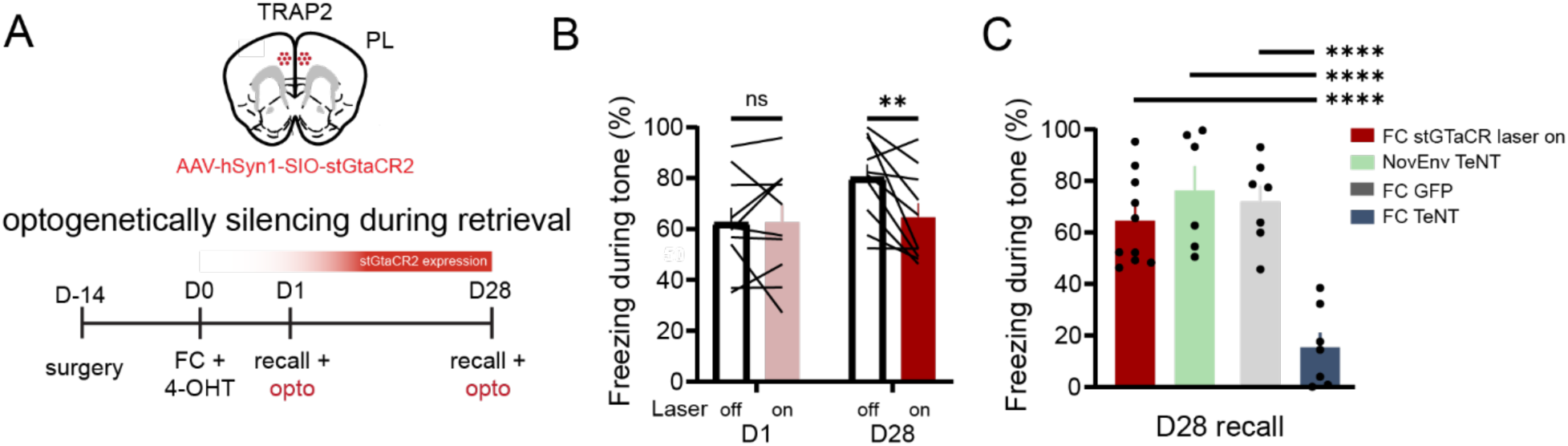
– Optogenetic silencing of learning-activated mPFC neurons mildly impairs remote memory. **A**. Schematic illustrating experimental strategy to optogenetically inhibit mPFC encoding ensemble during remote memory retrieval. B. Freezing during D1 and D28 memory retrieval in stGTaCR mice, comparing laser on and off epochs (Time F(1,18)=5.2, P=0.034; Laser F(1,18)=1.6, P=0.2; Time x Laser F(1,18)=5.6, P=0.028, Two-way ANOVA with Bonferroni’s multiple comparisons test, n=10 mice). C. Comparison of tone freezing during remote recall across groups (F(3,26)=17, P<0.0001, One-way ANOVA Tukey’s multiple comparisons test, n=10 stGTaCR mice, n=6 NovEnv-TeNT mice, n=7 FC-GFP mice, n=7 FC-TeNT mice). Data from NovEnv-TeNT, FC-GFP, and FC-TeNT groups are reproduced from main figure. **P<0.01, ****P<0.00001. Error bars represent mean ± s.e.m.

**Extended Data Figure 8.**
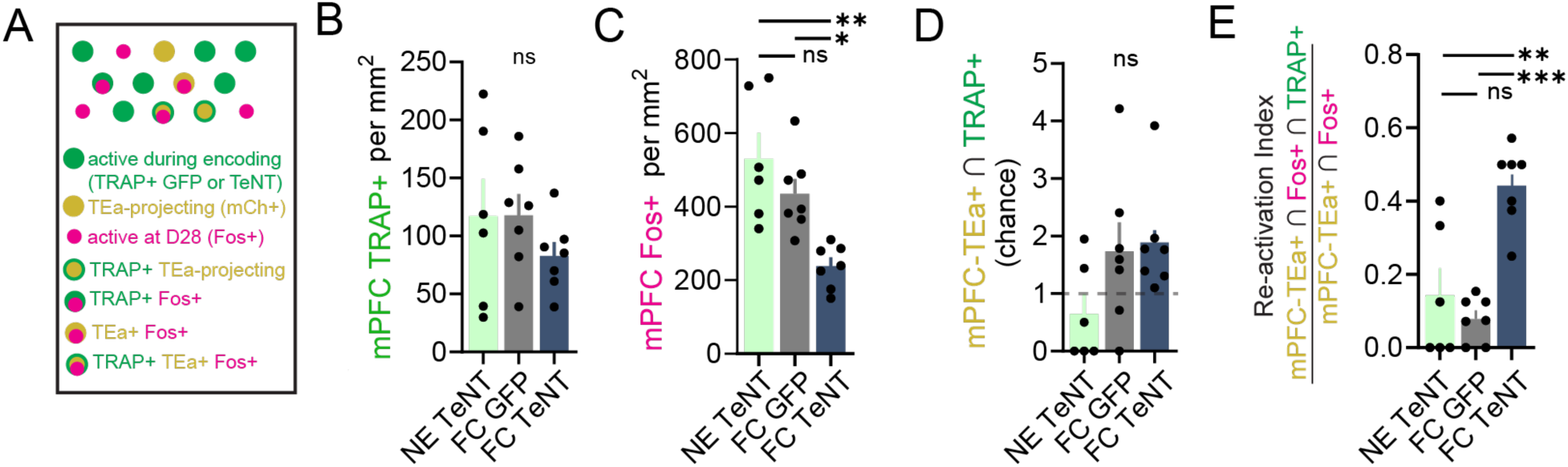
– Additional histological comparisons following synaptic silencing of mPFC learning ensembles. **A**. Schematic illustrating labeling strategy to examine remote ensemble membership. B. Density of mPFC neurons active during memory encoding (TRAP+) (F(2,17)=0.9, P=0.4, ANOVA, n=6 NE-TeNT mice, n=7 FC-GFP mice, n=7 FC-TeNT mice). C. Density of mPFC neurons active during remote recall (Fos+) (F(2,17)=7.3, P=0.005, ANOVA with Tukey’s multiple comparisons tests, n=6 NE-TeNT mice, n=7 FC-GFP mice, n=7 FC-TeNT mice). D. mPFC-TEa projecting neurons (mCh+) that were active during encoding (TRAP+), normalized to chance((F(2,17)=2.4, P=0.11, ANOVA with Tukey’s multiple comparison’s test; n=6 NE-TeNT mice, n=7 FC-GFP mice, n=7 FC-TeNT mice). **E.** Fraction of mPFC-TEa neurons (mCh+) active during recall (Fos+) and active during encoding (TRAP+) relative to total population of mPFC-TEA projecting neurons active during recall (F(2,17)=17.2, P<0.0001, ANOVA with Tukey’s multiple comparisons tests, n=6 NE –TeNT mice, n=7 FC-GFP mice, n=7 FC-TeNT mice). *P<0.05, **P<0.01, ***P<0.001. Error bars represent mean ± s.e.m.

